# Housing conditions affect adult zebrafish (*Danio rerio*) behavior but not their physiological status

**DOI:** 10.1101/2023.02.10.528020

**Authors:** Sara Jorge, Luís Félix, Benjamín Costas, Ana M Valentim

**Affiliations:** i3S - Instituto de Investigação e Inovação em Saúde, Universidade do Porto, Portugal; Laboratory Animal Science, IBMC - Instituto de Biologia Molecular Celular, Universidade do Porto, Porto, Portugal; Centro Interdisciplinar de Investigação Marinha e Ambiental, (CIIMAR), Matosinhos, Portugal; Instituto de Ciências Biomédicas Abel Salazar (ICBAS), Universidade do Porto, Porto, Portugal; Centre for the Research and Technology of Agro-Environmental and Biological Sciences (CITAB), University of Trás-os-Montes and Alto Douro (UTAD), Vila Real, Portugal; Instituto para a Inovação, Capacitação e Sustentabilidade da Produção Agroalimentar (Inov4Agro), UTAD, Quinta de Prados, 5000-801 Vila Real, Portugal

**Keywords:** zebrafish, environmental enrichment, cortisol, behavior, skin mucus, welfare

## Abstract

Zebrafish is a valuable model for neuroscience research, but the housing conditions to which it is daily exposed may be impairing its welfare status. The use of environmental enrichment and the refinement of methodology for cortisol measurement could reduce stress, improving its welfare and its suitability as an animal model used in stress research. Thus, this study aimed to evaluate (I) the influence of different housing conditions on zebrafish physiology and behavior and (II) skin mucus potential for cortisol measurement in adult zebrafish. For this, AB zebrafish were raised under barren or enriched (PVC pipes and gravel image) environmental conditions. After 6 months, the behavior was assessed by different behavioral paradigms (shoaling, white-black box test, and novel tank). The physiological response was also evaluated through cortisol levels (whole-body homogenates and skin mucus) and brain oxidative stress markers. Results revealed that enriched-housed fish had an increased nearest neighbors’ distance and reduced activity. However, no effect on body length or stress biomarkers was observed; whole-body and skin mucus cortisol levels had the same profile between groups. In conclusion, this study highlights the skin mucus potential as a matrix for cortisol quantification and how environmental enrichment could influence the data in future studies.

**Simple Summary:** The delivery of proper housing conditions may translate into good fish welfare. As zebrafish housing is usually poorly enriched, the fish could be unable to express some natural behaviors, leading to distress and/ or stress mechanisms’ dysregulation. This work focused on the examination of zebrafish welfare raised under different housing conditions (barren or environmentally enriched) and the testing of a low-invasive technique (skin mucus collection) to measure the main stress hormone (cortisol). The data was processed to assess body length, behavior, and physiological status. Results revealed that enrichment induced minor alterations on zebrafish behavior. Thus, the influence of housing conditions should be considered in future research, depending on the purpose of the study. Also, skin mucus appears to be a promising matrix to replace whole-body to measure cortisol in zebrafish, since its collection is nonlethal and showed similar results to the traditional method.

## 1. Introduction

Over the years, the scientific community has been raising and housing zebrafish in barren tanks to reduce experimental variation, facilitate health monitoring and husbandry practices. However, in the wild, these fish encounter enriched environments [1] and a variety of stimuli that have a direct impact on their behavioral and physiological responses. Therefore, the exposure to laboratory conditions not only increases the expression of stereotypical behaviors and stress but also limits captive animals from expressing a complete behavior repertoire [2]. Nevertheless, zebrafish use as a laboratory model in neuroscience continues to rise [3] and efforts have been made by the scientific community to ameliorate the deleterious effects of captivity.

Enrichment aims to enhance laboratory animals’ welfare by changing their housing environment according to their natural history and allowing the expression of species-specific behaviors [4]. To date, zebrafish showed a preference for environmental enrichment over barren environments [5,6], such as gravel, gravel images, or plants [5]. In addition, when an unpredictable chronic stress was used, the enrichment (gravel, plants, plastic structure) attenuated its effects on behavior and physiological parameters, reducing animals’ vulnerability to stress [4].

Overall, pipes and plants have been used in fish species to provide shelter [7] against aggressive conspecifics and tank disturbances [8,9]. Despite this, the literature describes opposing effects when plants are employed alone or in combination with other enrichment forms in zebrafish housing (e.g., increased [10] or reduced [11,12] aggression). Furthermore, the toxicological and biological concerns (e.g., plastic microparticles and/or substances; biofilm development) [13] hampered its application; the same happens with gravel use or another substrate that requires constant cleaning to avoid biofilm creation.

Alternatively, gravel images are available for commercial use [14] and do not interfere with husbandry procedures or water quality, but their beneficial effect on zebrafish welfare is still unclear. For instance, enrichment of a gravel image and a floating plant for 4 months did not induce an effect on zebrafish stress recovery [15]. Thus, clarification is needed regarding the best choice of enrichment for zebrafish and to which extent should be used.

Here, the main aim of this study was to assess the influence of different housing conditions (barren versus enriched with PVC pipes and gravel image) on zebrafish body length, physiology, and behavior. The combination of two enrichments such as gravel images and pipes are appealing for the zebrafish welfare improvement since it could increase the animal sense of safety by providing hiding spaces/ shelters, and background for camouflage against endangering situations, favoring the expression of natural behaviors. Therefore, it was hypothesized that distress, anxiety-like behaviors, and abnormal behaviors would be attenuated under environmental enriched conditions.

As behavioral and physiological analysis are commonly used to study distress [16], the secondary aim of this work was to test skin mucus collection as a new methodology for cortisol measurement in zebrafish. Cortisol is usually extracted from whole-body homogenates in zebrafish, which implies the animals’ death and hampers the possibility of repeated measurements [17]. Skin mucus is a promising matrix to be studied because its composition has been described to be altered by the fishes health in a manner comparable to plasma; in addition is a non-terminal collection, minimally invasive, practical [18], and quick to collect. At the end, similar outcomes in both cortisol matrices should be expected, supporting skin mucus as a less intrusive matrix.

## 2. Materials and Methods

### 2.1. Ethics statement

All experiments were approved by the Animal Welfare and Ethics Review Body of the i3S (2021-21) and conducted by researchers with personal licenses to work with animals approved by the National Competent Authority for animal research (Direção-Geral de Alimentação e Veterinária) and in agreement with European Directive and Portuguese legislation (2010/63/EU and 113/2013, respectively) on the protection of animals used for scientific purposes.

### 2.2. Animals and housing

Wild-type AB zebrafish were bred in-house. The embryos were disinfected with Chloramine-T 0.5% solution of 0.0037%, before being housed in groups in glass tanks (24 × 45 × 25 cm) of 20L capacity. All tanks were part of an open system supplied with UV-sterilized tap water (pH 7.1 ± 0.3), maintained under controlled temperature (28.3 ± 0.4 °C) and photoperiod (14:10 h light: dark). Tanks were placed on top of white styrofoam and illuminated by ceiling-mounted light emitting diodes. Zebrafish were hand fed twice a day with ZEBRAFEED for larvae, juvenile and adults (<100, 100-200, and 400-600 μM, respectively; Sparos Lda, Olhão, Portugal) according to the manufacturer’s instructions. Feed deprivation was applied 24 h before euthanasia to discard the influence of feeding on cortisol levels. The tanks maintenance was done when necessary, without changing the location of structures and equipment.

### 2.3. Experimental design

After embryos disinfection, 150 embryos were placed in each study tank, where they hatched and were maintained up to 6 months in the same housing conditions – barren or enriched; thus, the tank was the experimental unit (n= 5). Two methodological replicates, i.e., two batches of animals were used. Each batch was randomly distributed into the housing conditions; the first replicate comprised two tanks for each condition and the second replicate three tanks per condition, and this was taken in consideration in the statistical model. In both conditions, the tanks were equipped with a heater and thermometer. The enriched tanks included a gravel image (Figure S1) placed in the exterior bottom of the tank, and 3 PVC pipes sections (6.1 cm length; 3 cm external diameter; 2.5 cm internal diameter) with a dark grey color (Figure 1).

**Figure 1.**
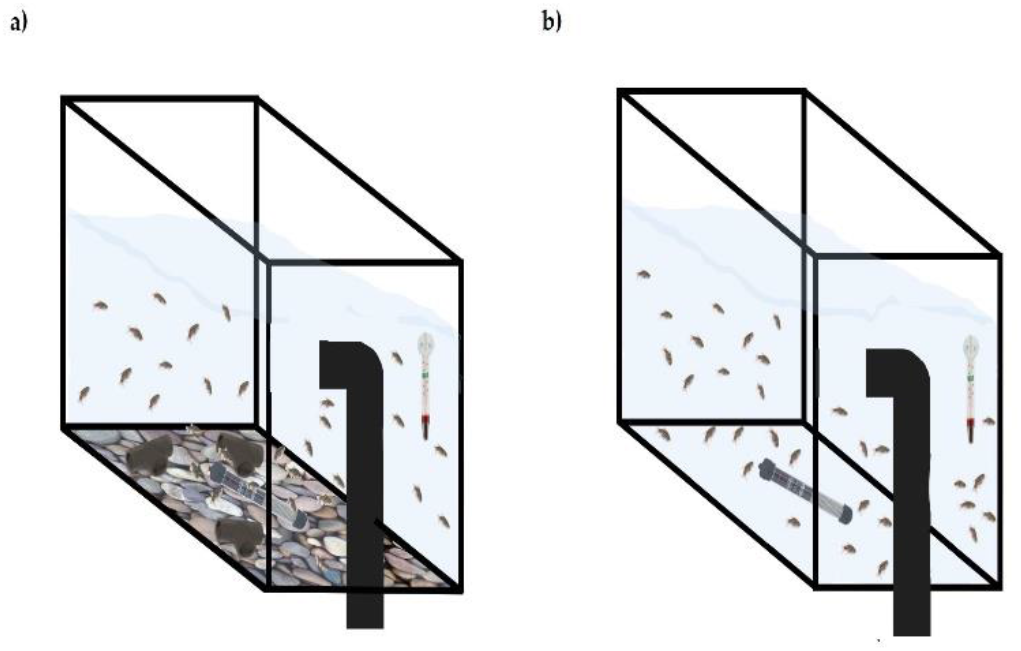
Schematic representation of enriched (with 3 PVC pipes and gravel image); (**a**) and barren (**b**) housing. All tanks had a heater and a thermometer inside the tank.

The rationale for using pipes and gravel images in this study was to give the animals a sense of safety through the availability of shelters (pipes) and camouflage (gravel image) against potential threats. The gravel images were also chosen because of the Schroeder, et al. [5] study of preference, which demonstrated the zebrafish preference for this enrichment type over barren conditions. Moreover, both enrichment items are affordable and easily maintained, reducing the workload and expenses in the research facilities. The items also do not represent a risk for the water quality as the plumbing pipes are inert and the images were placed outside the tank.

After 6 months in these conditions, the animals were subjected to behavioral recordings and sampling.

### 2.4. Behavioral testing

The behavioral tests took place in the same room where the animals were housed one day before the animals were sampled for biochemical analysis. The recordings were made with a digital video camera and started at 10:00 am and ended at 16:00 pm with the following order: shoaling, white/black tank and novel tank test, involving different animals in each test. The water of each behavioral apparatus was fully replaced by fresh system water between animals/ trials. To prevent sampling the same fish twice, fish were transferred to another tank after testing. The animals were also randomly distributed between tests. A researcher was blinded to the housing condition and batch when analyzing the behavioral data.

Before behavioral testing, some home-tank recordings of three enriched tanks (Video S1) were also performed for three days (two times a day; morning and afternoon) to observe the number of entries and exits in the front pipe (the most visible one) and conclude if the animals use or interact with these structures.

#### 2.4.1. Shoaling test

Because zebrafish shoals tighten in stressful or threatening events [19], the shoaling test was used to evaluate group cohesion. For this, five fish per tank were placed in a 24 × 24 cm glass tank with a water column of 4 cm. After 30 min of habituation to the surroundings, the animals’ shoaling behavior was video recorded from above for 10 min. The software The Real Fish Tracker [20] was used with a confidence threshold of 40, a mean filter size of 3 pixels to quantify the average inter-fish distance and the neighbor nearest (IDNN) and farthest (IDFN) distance (cm).

#### 2.4.2. White/black tank test

The white/black tank test is an anxiety test, based on zebrafish natural preference for dark backgrounds over white and bright environments [21]. The test was conducted in a tank (20 × 10 cm) divided evenly between a black and white side and filled with a 4 cm water column. A total of 25 fish per housing condition (5 fish tested per tank) were analyzed. Each fish was placed on the white side of the tank and its behavior was recorded for 7 min. Time (s), distance traveled (m), average speed (m/s), immobility (s), and the number of entries on the white side were measured. The latency to enter on the black side and re-enter on the white side were also determined. The behavioral endpoints were obtained using the Any-mazeTM behavioral tracking software (Stöelting, Dublin, Ireland).

#### 2.4.3. Novel tank test

The novel tank test considers new environments to be anxiogenic for zebrafish, inducing higher occupation of the tank bottom and then progressive habituation to the environment with increased exploration at the top of the tank [22]. Hence, in this study, the animals (25 fish per housing condition; 5 fish collected per tank) were individually transferred to a new tank (24 × 12 × 8 cm) filled with a 12 cm water column and space occupation and exploratory behavior video recorded for 6 min using a top camera. The tank was virtually divided into an upper (UP) and bottom (BTM) zone. The distance traveled (m), average speed (m/s), angular velocity (°/ s), time (s), immobility (s), and erratic movements were analyzed using the Any-mazeTM software as previously described in Jorge, et al. [23].

During the observation of the novel tank videos, a repetitive behavior was noted, whereat zebrafish touched the tank walls with its mouth several times using a circling behavior. To ensure this would not interfere with the observation of the other variables tested in the novel tank, this repetitive behavior was quantified as the average of wall contacts (frequency and duration) using the BORIS v. 7.13.5 (Behavioral Observation Research Interactive Software; Friard and Gamba [24]) software. Each contact started after the fish touched the tank wall with its mouth for the 4th time and ended with the last contact in the wall before the fish swims in the opposite direction.

### 2.5. Sample collection

The day after behavioral testing, three fish per tank were euthanized with MS222 (250 mg/L). Then, two fish were placed on a sponge soaked with euthanasia solution, and sterile swabs (155C, Copan, Brescia, Italy) were used to swab the left flank of each fish six times, from the pectoral fins to the beginning of the caudal fin. A swab rotation was done in the middle of each swabbing as an attempt to maximize mucus collection. The cotton tip of the two swabs were then pooled in an Eppendorf tube with 500 μL ice-cold PBS (phosphate buffered saline, pH 7.4) to obtain a meaningful cortisol value for the assay analysis.

Next, the head of the third fish was discarded, and the trunk collected in 5mL of ice-cold PBS. Males were immediately sampled, whereas females had their eggs extracted [25] and their trunks washed with ice-cold PBS prior to its collection, to detach any eggs and avoid cross-contamination between cortisol matrices. The sample collection was repeated until 4 samples per matrix in each tank were obtained. The dissection material was disinfected with 70% alcohol and cleaned with ice-cold PBS between each animal. The Digimizer, MedCalc Software (Version 5.3.5, Ostend, Belgium) was used to analyze photos of eight fish per tank acquired during sampling to assess how housing affects growth. Total body length was measured as the distance between the most anterior section of the fish head and the caudal fin end. Additionally, from the fish collected to cortisol analysis, six brains were randomly chosen from each tank for the oxidative stress analysis. Each sample was collected in less than 1 min and stored at -20 ºC until processing. These animals were not tested before in the behavioral tests described.

### 2.6. Biochemical analysis

One pool of six zebrafish brains per tank was homogenized in ice-cold buffer (0.32 mM of sucrose, 20 mM of HEPES, 1 mM of MgCl, and 0.5 mM of phenylmethyl sufonylfl uoride, pH 7.4) [26] by bead beating in a Tissuelyser II (Quiagen, Hilden Germany; 30 sec at 30 Hz; one 4.5 mm steel bead/ sample). Following homogenization, the samples were centrifuged at 15000xg for 20 min in a cooled centrifuge (4 °C; Prism R, Labnet International Prism-R, Edison, USA) and their supernatant was collected for measurement of oxidative stress biomarkers. Hence, the reactive oxygen species (ROS) were measured at 480 (excitation) and 530 (emission) nm according to [26,27], using 2′,7′-dichlorofluorescein diacetate as probe dye. The superoxide dismutase (SOD) and catalase (CAT) activities were determined by nitroblue tetrazolium (NBT) reduction at 560 nm [28] and hydrogen peroxide at 240 nm [29], respectively. The conjugation of 1-chloro-2,4-dinitrobenzene with reduced glutathione (GSH) at 340 nm was used to measure the glutathione-s-transferase (GST) activity. The glutathione reductase (GR) and glutathione peroxidase (GPx) activities were measured by the oxidation and reduction of NADPH at 340 nm as described in Massarsky, et al. [30]. The GSH and the oxidized glutathione (GSSG) states were quantified at 320 nm (excitation) and 420 (emission) nm according to Gartaganis, et al. [31]. The oxidative stress index (OSI) was given by the GSH:GSSG ratio. The thiobarbituric acid reactive substances (TBARS) activity reflected the degree of lipid peroxidation and was measured at 530 (MDA-TBA adducts and 600 (non-specific adducts) nm. Carbonyls (CO), the protein oxidation indicators, were determined through the DNPH (2,4-dinitrophenylhydrazine) method of Mesquita, et al. [32] at 450 nm.

The acetylcholinesterase (AChE) activity was analyzed at 405 nm on microplates [33], based on the Ellman’s method [34]. The method of Domingues, et al. [35] was used to assess the activity of lactate dehydrogenase (LDH) at 340 nm.

All samples were run in duplicate and measured against a reagent blank at 30 °C using a PowerWave XS2 microplate scanning spectrophotometer (Bio-Tek Instruments, USA) or a Varian Cary Eclipse (Varian, USA) spectrofluorometer. The protein content within each sample was determined at 280 nm in a BioTek Take3 microvolume plate (Bio-Tek Instruments Inc., Winooski, VT, USA).

### 2.7. Cortisol extraction and analysis

Zebrafish trunks were cut with an ophthalmologic scissor (Dentalhonest, Sichuan, China) in 500 μL PBS, before being homogenized by bead beating (five 3 mm steal beads) in the FastPrep®-24 (MP Biomedicals, Solon, USA; 6 m/s for 60 s) at room temperature. Skin mucus samples were only vortexed (2 min; 35 Hz) before solvent extraction. Afterward, 500 or 750 μL methanol (HPLC grade ≥99.8%; Fisher Scientific, Loughborough, UK) was added to each sample of mucus or trunk, respectively. Then, the samples were placed overnight for 24 h at room temperature in a lab roller (60 rpm), built in-house according to Dhankani and Pearce [36]. On the next day, the samples were 10000xg centrifuged for 10 min in a cooled centrifuge (4°C; Centrifuge 5415 R, Eppendorf, Hamburg, Germany), and the supernatant transferred to new microcentrifuge tubes, before being placed on a vacuum concentrator (Savant™ SPD131DDA SpeedVac™ Concentrator, ThermoScientific Inc., Waltham, USA) at 36 ºC; the swabs were removed from the mucus samples tubes before evaporation. Following the solvent evaporation, 500 or 125 μL of assay diluent was added to each trunk or mucus sample, respectively. Then, the samples were incubated overnight in a refrigerator at 4 ºC. On the following day, 500 μL n-Hexane (97+ %; Acros Organics, Geel, Belgium) was added to trunk samples to remove the interference of precipitated lipids. These samples were then frozen at -20 ºC for 15 min, followed by the removal of the organic layer. After this step, all samples were analyzed according to the instructions of the ELISA kit (Salimetrics® Cortisol Enzyme Immunoassay Kit; #1-3002, Salimetrics, State College, PA, USA).

Following the assay, the samples protein content was measured at 280 nm using NanoDrop One (NanoDrop Ins., Thermo Fisher Scientific, Waltham, MA, USA). The cortisol data are expressed in pg/mg protein.

### 2.8. Statistical analysis

The data normality and homogeneity were confirmed by Shapiro-Wilk and Levene’s tests, respectively. Whenever necessary the data were log-transformed to achieve normality. Significant differences were considered at *p* <0.05 (two-tailed). Graphical representations were plotted in GraphPad Prism 7 for Windows (GraphPad, Inc., San Diego, CA).

Data generated were analyzed using the IBM SPSS Statistics 27.0 computer program (SPSS, Chicago, IL, USA). As the experimental unit was the tank, an average of the variables’ values corresponding to the animals from the same tank was used for the statistical analysis. The housing condition and batch were considered fixed and random factors, respectively. Univariate analysis of variance or Mann-Whitney U-test were used to assess differences between groups, whereas Student’s paired t-test or Wilcoxon signed rank test were used to determine differences within groups. For the novel tank and white/black tank data, the one-sample t-test was used to compare the time spent in each zone of the tank to the time that would be obtained by chance (180 and 210 sec, respectively). One animal from the barren housing was excluded from the novel tank analysis due to poor video quality.

## 3. Results

The fish length was not significantly influenced by either housing or batch. The animals from the barren housing had an average length of 3.80 ± 0.18 cm, whereas the enriched-housed fish had an average of 4.15 ± 0.36 cm at 6 months of age.

The number of entries and/ or exits (total of 32 passages/ min) of animals in the front pipe from the enriched environment proved that the animals did use the pipes (Video S1).

### 3.1. Shoaling

In the shoaling test, it was shown that the housing conditions did not influence the average inter-fish distance (Figure 2a) and the IDFN (Figure 2b). Nonetheless, the shoals from enriched tanks had a significantly higher IDNN (Figure 2c; *p*=0.031) compared to the shoals from barren housing. Also, no batch interaction was noted.

**Figure 2.**
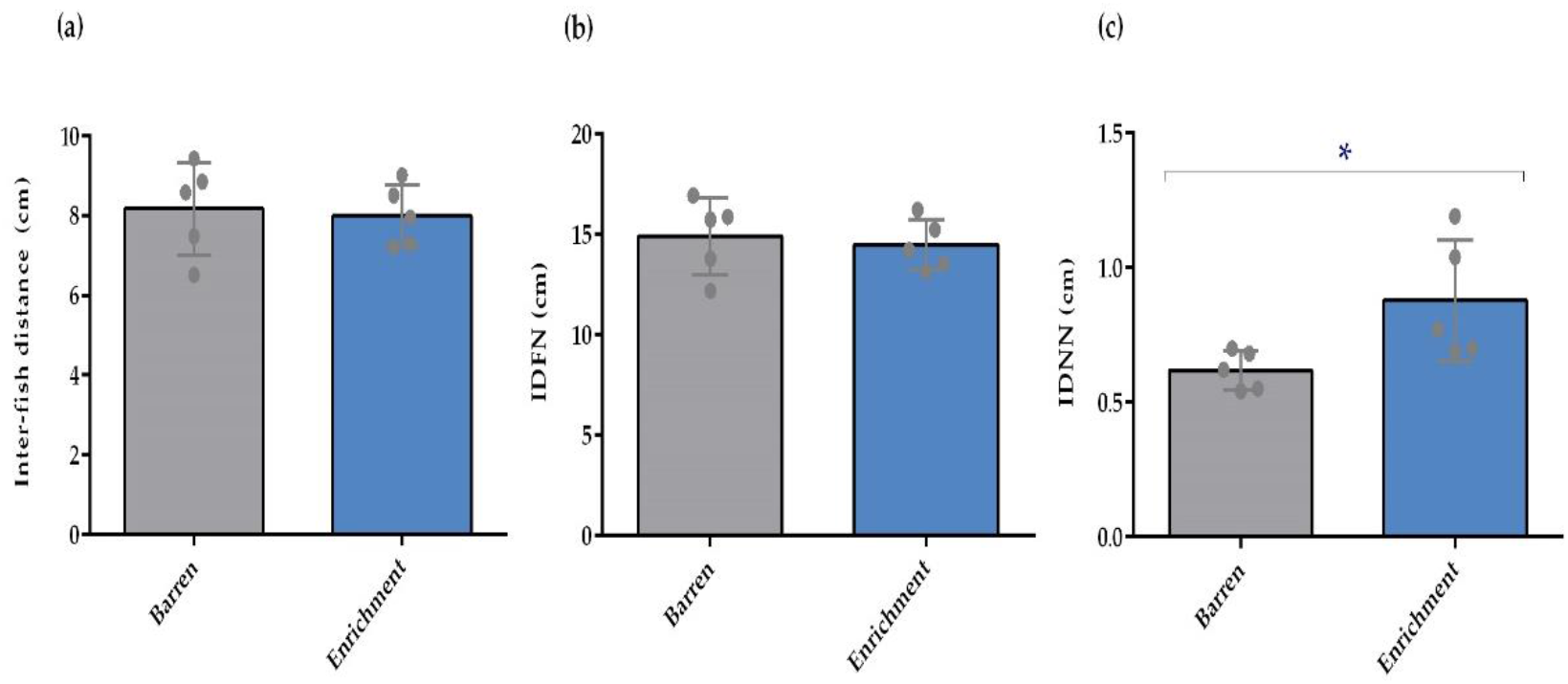
Shoaling behavior of adult zebrafish after exposure to barren and enriched conditions (n=5). (**a**) Individual inter-fish distance (cm); (**b**) Distance of the farthest neighbor (IDFN; cm); (**c**) Distance of the nearest neighbor (IDNN; cm) of enriched and barren housed fish. Data are expressed as mean ± standard deviation. \**p* < 0.05 for comparison between housing conditions. Each point represents one experimental unit, i.e., a tank.

### 3.2. White/black tank

In the white/black tank test (Figure 3), both barren and enriched conditions significantly increased the time spent on the black side compared to the white side (*p*= 0.043 and *p*<0.001, respectively).

**Figure 3.**
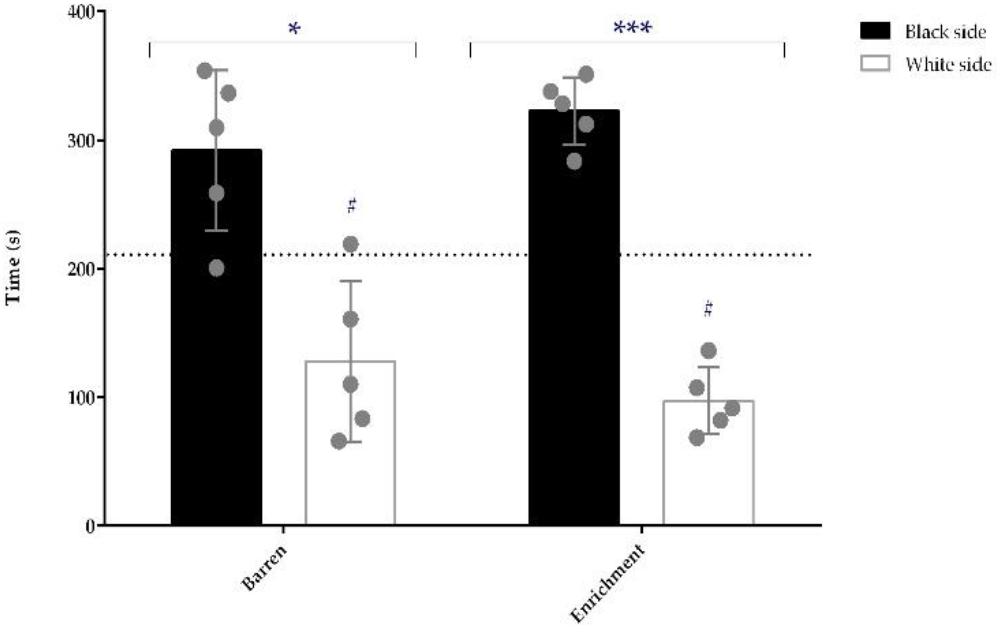
Time spent (s) in each side of the white/black tank by enriched and barren-housed adult zebrafish (n=5). Data are presented as mean ± standard deviation. \**p* < 0.05 and \*\*\**p* < 0.001 for comparisons between the white and black side of the tank for the barren and enriched-housed animals, respectively; # *p* < 0.05 for comparison between the time spent in the white side and the time spent there by chance (210 s). Each point represents one experimental unit, i.e., a tank.

In addition, fish from both conditions spent significantly less time (*p*= 0.042 and *p*<0.001 for barren and enriched conditions, respectively) on the white side compared with the value predicted by chance (210 sec). No differences between groups, batch or interaction between batch and treatment were detected.

### 3.3. Novel tank

The novel tank analysis revealed that animals housed in barren tanks swam more than the ones housed in the enriched environment (*p*= 0.009; Figure 4a). These animals spent more time in the BTM compared to the UP zone of the tank (*p*=0.042) whereas barren tanks did not induce different space occupation (Figure 4b). Supporting this, the enriched-housed fish spent less time in the UP zone when compared with the value that would be expected by chance (180 sec; *p*=0.042; Figure 4c).

**Figure 4.**
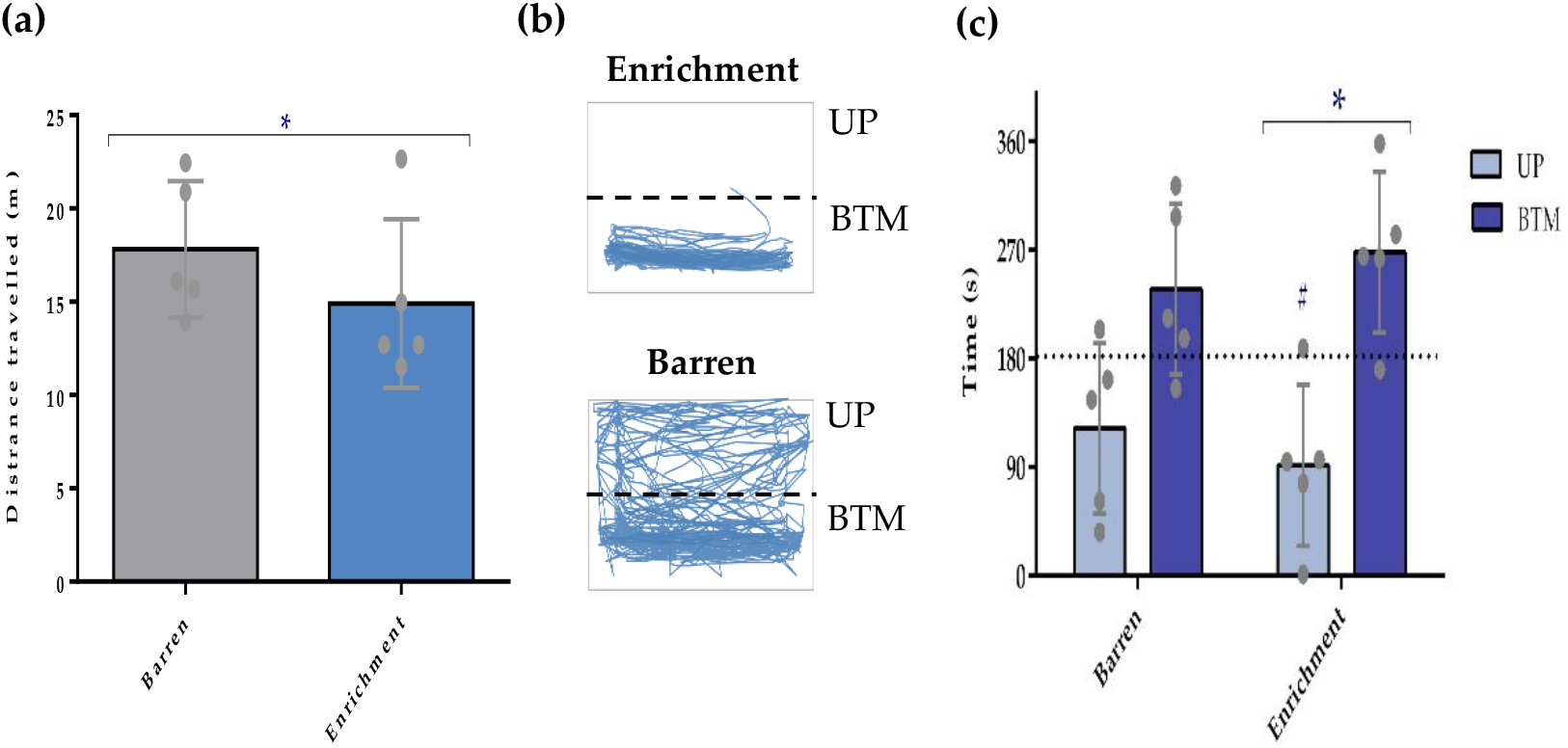
Locomotor activity of adult zebrafish during the novel tank test (n=5). (**a**) Total distance travelled (m) by barren and enriched housed fish; (**b**) Representative tracking example of one fish from each housing condition showing different behavioral patterns (AnyMaze software); (**c**) Time spent (s) in each zone (UP and BTM) per housing condition. UP-upper zone of the tank; BTM-bottom zone of the tank. Data are expressed as mean ± standard deviation. \**p* < 0.05 for comparison between housing conditions in (a); \**p* < 0.05 for comparison between UP and BTM in enriched-housed animals in (c); # *p* < 0.05 for comparison between the time spent in the UP zone and the time spent there by chance (180s). Each point represents one experimental unit, i.e., a tank.

Nevertheless, there were no other differences detected between treatment groups, nor interactions between treatment*batch. However, a batch effect on the locomotory activity and on high-speed movements (distance and duration swam) was observed (*p*<0.05) (Table S1).

From the 25 animals observed in the novel tank per condition, two from the barren tank and four from the enriched tank did not present the repetitive behavior previously described. Also, no significant effects of housing or batch were detected in the duration and frequency of this behavior.

### 3.4. Biochemical analysis

No significant effect (Figure S2) of housing or batch was observed in the oxidative stress biomarkers nor in the AChE and LDH activity.

### 3.5. Cortisol analysis

There was no difference between treatments (housing conditions), nor a significant effect of batch on cortisol levels (Figure 5) measured in the whole-body or in the skin mucus of the animals.

**Figure 5.**
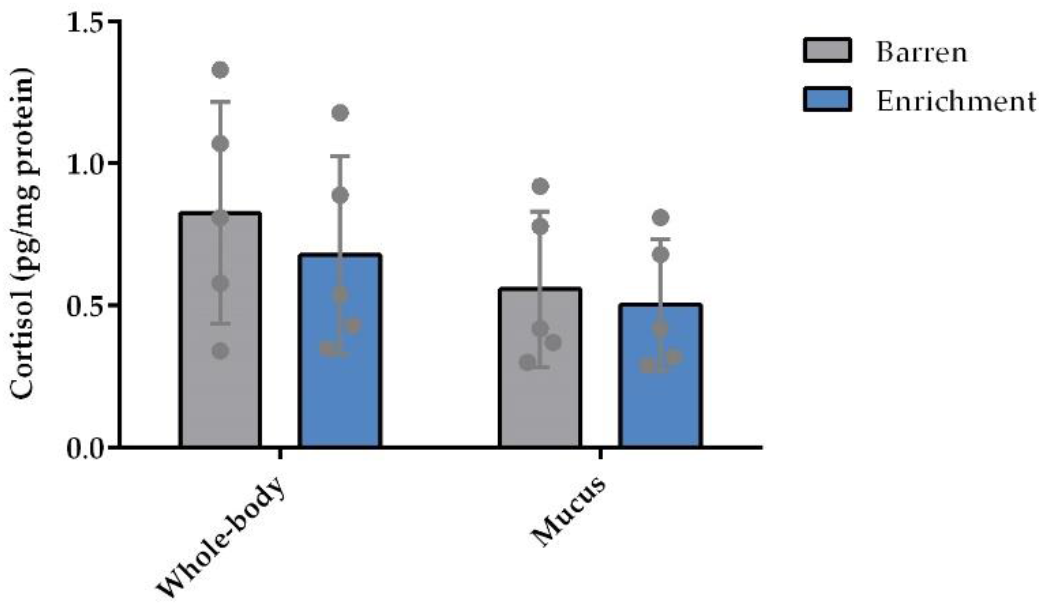
Whole-body homogenates and skin-mucus cortisol levels (pg/mg protein) after a 6-months exposure to enriched and barren housing (n=5). Data represented as mean ± standard deviation. Each point represents one experimental unit, i.e., a tank.

## 4. Discussion

Inadequate housing may affect zebrafish welfare and induce behavioral and physiological changes, which can impact the research outcomes [37]. In this study, the influence of different housing conditions on body length, behavior and physiology of zebrafish was evaluated, and it was concluded that, depending on the parameter examined, environmental enrichment may influence animal behavior, but it was unclear whether the enrichment used in this study would be favorable for zebrafish.

Firstly, no effects on fish total body length were observed after exposure to different housing conditions. Another study [38] showed an increase in the zebrafish length housed in environmental enrichment, but only at 60 dpf, while no differences were detected at older stages; thus, it may be possible that the influence of housing conditions on growth is only detected at earlier stages.

Regarding behavior, the different housing conditions (enriched vs barren) elicited a minor alteration on the shoaling test. Fish raised in barren tanks had a lower IDNN than fish raised in enriched ones, even though the average inter-fish distance and IDFN were not altered between treatments. Nonetheless, Wilkes, et al. [39] suggested that when zebrafish are introduced to a novel environment, an initial cohesiveness may occur in less than 24 hours. Therefore, it is possible that this IDNN was a natural response induced by novelty and would dissipate throughout time. A high cohesion may be an anti-predator defense mechanism in a new environment [40], but has also been described to increase when free-choice exploration was given [41]; thus, this behavior presents an ambivalent valence, hard to interpret. Nevertheless, the measure that gives us the overall shoaling cohesion is the inter-individual distance [42], which was not altered.

To clarify if stress and/ or anxiety were altered, the animals were tested in two classical anxiety tests. In the white/black tank test there were no differences between groups in the several variables studied. All animals spent more time in the black than in the white side, as expected for a control animal. This can be interpreted as an avoidance indicator [43,44] rather than anxiety since these animals naturally prefer darker backgrounds that allow them to be safely hidden. Manuel, et al. [45] described that enriched-raised zebrafish were less anxious in this behavioral test than the animals raised in a barren environment. However, the strain used was different, which could influence the outcomes of the results [46]. In addition, we analyzed the test continuously, while Manuel and colleagues analyzed minute by minute; the differences reported between housing conditions were only detected in minutes 2, 3, and 4, indicating that barren-raised animals rapidly behaved as the enriched-raised animals. In our study and in the referred study, animals raised on both housing conditions spent more time in the black than in the white side of the tank, showing their natural preference.

The other classical anxiety test, novel tank, revealed one difference between groups. The fish raised in barren conditions swam more than the ones raised in enriched environments. Although an increase in fish locomotion may indicate anxiety [47], the proportion of the increase is often compared with a baseline. In the scenario of different housing conditions, a baseline is hard to define. Nevertheless, there were no differences between treatments regarding time and distance swam at high speed (higher than 0.05–0.07m/s); these movements at high speed are often associated with erratic movements, a stress-related behavior [48]. Thus, the increase in distance swam by the fish raised in barren conditions in the novel tank test may be due to the environment’s resemblance to their original housing, which could have caused rapid habituation to the new settings and an increase in the exploratory activity; whereas the enriched-raised fish were used to have shelters available and a tank with a different background compared to barren housing and novel tank environment. This is further supported by the lack of differences in space occupation in the barren-treated animals, while the enriched-treated animals spent more time in a more protected zone, the bottom of the tank. Hence, the fish from barren housing might be bolder to explore the novel tank, whereas enriched-reared fish were more reluctant regarding its exploration [49].

Previous work [50,51] demonstrated that three or seven days of unpredictable chronic stress in zebrafish larvae were enough to build stress resilience in adults, inducing no alterations or decreased anxiety, respectively. The barren rearing conditions are very different from the natural ones [52], where larvae have shelters available (rocks, plants) and different background colors to choose, depending on the type of sand and gravel. Thus, raising zebrafish in barren conditions may lead to early-life stress, not responding with the expected stress to a novel environment such as the novel tank. However, we can only speculate, as we did not use a stress protocol, but only placed the animals alone in a novel tank, which may be not stressful enough to trigger a stress resilience response.

Stress resilience was also shown when environmental enrichment (plants, shelters, and gravel) was introduced to adult zebrafish for several days (15-28): chronic [4] and acute [53] stress did not elicit behavioral or physiological alterations in the enriched-housed animals compared with barren-housed animals. However, in another study [15], enrichment (gravel images and plants) did not induce a recovery after simulated predator presentation, air emersion, or fin clipping. Thus, the type of stressors and/ or the type of environmental enrichment (e.g., having gravel vs gravel images) may be crucial to elicit stress resilience, but more research needs to be done to answer these issues.

In the novel tank analysis, the detected repetitive behavior towards the wall of the tank may be related to the material of the tank that reflects the animals’ image and not by the housing condition per se, as the behavior was equally frequent in both groups.

Although small behavioral alterations were observed, there were no differences in the physiological measures; cortisol levels and the brain biochemical analysis were similar between housing conditions. Contrary to our study, Marcon, et al. [55] showed a decrease in ROS levels and an increase in catalase (antioxidant) activity in enriched environments compared to barren conditions, which may confer a protection against stressful events. Nevertheless, there is a difference between this study and ours that could have contributed to the observed outcomes: the timing to introduce environmental enrichment; our animals were raised in different environments, while, in the referred study, adult zebrafish were placed for 21 and 28 days in an enriched environment.

Cortisol is the main stress biomarker in fish, but in this study, animals housed in different conditions had similar cortisol levels. This may occur because the animals were housed in these specific housing conditions from fertilization and only experienced modest husbandry procedures (e.g., debris removal) for months, thus the animals may have adapted to the conditions and the animal facility routine. If a stress protocol would be applied, the animals could respond differently with distinct cortisol levels depending on the housing condition, as previously shown [4,53]. Here, enrichment may have dampened the zebrafish cortisol response when animals were subjected to unpredictable chronic or acute stress. Nevertheless, both studies share the same result as ours when the animals were housed in different conditions but not subjected to any stress protocol. Therefore, to corroborate the existing literature, future research using the enrichment employed in our study should include stressors testing as additional treatment groups.

Recognizing that depending on how much glucocorticoids (e.g., cortisol) the fish are exposed to during a stress response, its redox state can be changed [56] by heightened oxidative damage or antioxidant defenses [57]. If enriched or barren environments functioned as a stressor, it would be expected that the fish brains present an altered enzymatic profile (e.g. [4,55]). The stability of antioxidant defenses across different housing situations observed in our study indicates that fish raised in different housing conditions had similar stress biomarkers levels in adults, when no stress protocol is applied.

As cortisol is an important welfare/ stress indicator, its measurement should be practical and without side effects for the animal and data quality. Thus, we also tested a low-invasive and non-terminal method, skin mucus, that showed a similar cortisol profile between groups compared with the standard methodology, the whole-body. This approach will allow the reduction of the animals used in an experimental setting, as the animals can be re-used, for example for breeding purposes.

## 5. Conclusions

In summary, findings from this study demonstrate how housing enrichment from fertilization might influence adult zebrafish welfare and the practicability of using skin mucus for cortisol measurement. Overall, the use of the proposed enrichment (shelter and gravel image) did not interfere with the physiological stress (cortisol and redox status), nor with the anxiety test black/white tank. However, a minor alteration was detected in the shoaling test and the housing conditions interfered with the distance swam, and the anxiety-like profile in the novel tank test.

Given that fish may respond differently depending on the fish strain, enrichment duration, timing (ontogeny), and type, it is essential that the upcoming literature fully describes these crucial aspects. Also, the use of a stress protocol is useful to clarify how fish in different housing conditions respond to a stressor and how the hypothalamic-pituitary-interrenal axis is affected, demonstrating the real impact of housing on zebrafish welfare. Lastly, the similar cortisol levels profile of skin mucus and whole-body strengthens the potential to use skin mucus to measure cortisol as a non-terminal methodology.

## Supplementary Materials

The following are available online at www.mdpi.com/xxx/s1, Figure S1: Representation of the gravel image used on the tank bottom at the enriched housing; Figure S2: Status of the biochemical parameters measured in brain of the adult zebrafish housed in enriched and barren environments for 6 months. AChE - Acetylcholinesterase; CAT - Catalase; SOD - Superoxide dismutase; GPx - Glutathione peroxidase; GR - Glutathione reductase; GSH - Reduced glutathione; GSSG - Oxidized glutathione; GST - Glutathione-s-transferase; LDH - Lactate dehydrogenase; OSI - Oxidative stress index; ROS - Reactive oxygen species; TBARS - Thiobarbituric acid reactive substances. Data represented as mean ± standard deviation, except CAT and OSI, which data are presented as median [IQR]; Table S1: Significant data (p < 0.05) of the random factor Batch from the novel tank test that are not represented in the text or figure.; Video S1: Example of a brief enriched home-tank recording to show the animals’ interactions with the pipes.

## Author Contributions

Conceptualization, AMV and LF.; methodology, SJ, LF and AMV.; investigation, SJ and LF.; formal analysis and writing- original draft preparation, SJ.; supervision, AMV.; writing-review and editing, AMV, LF, BC and SJ. All authors have read and agreed to the published version of the manuscript.

## Institutional Review Board Statement

All procedures were approved by the National Competent Authority for animal research (Direção-Geral de Alimentação e Veterinária, Lisbon, Portugal) and by the Animal Welfare and Ethics Review Body of the Institute for Research and Innovation in Health (i3S).

## Acknowledgments

The authors would like to thank to the fish facility staff and to Raquel Vieira for the technical assistance in the stress oxidative assessments.

