## Supplemental tables and files for "Housing conditions affect adult zebrafish (*Danio rerio*) behavior but not their physiological status"

**Figure S1**

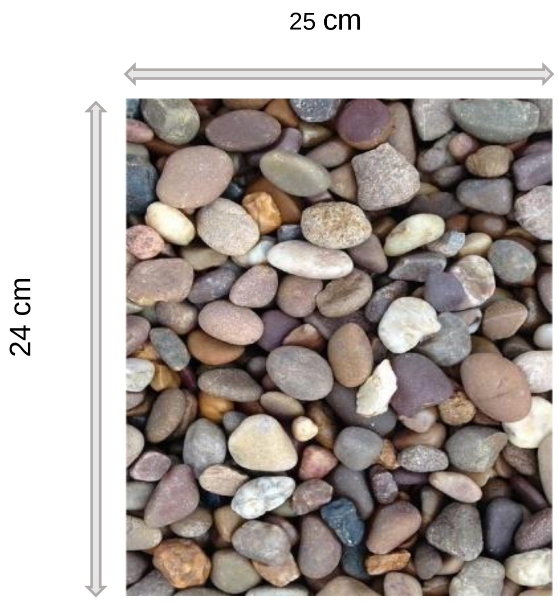


Source: https://pixabay.com

Reference:
20QTZ3-1000

The image was free for commercial use and no attribution required

**Figure S2**

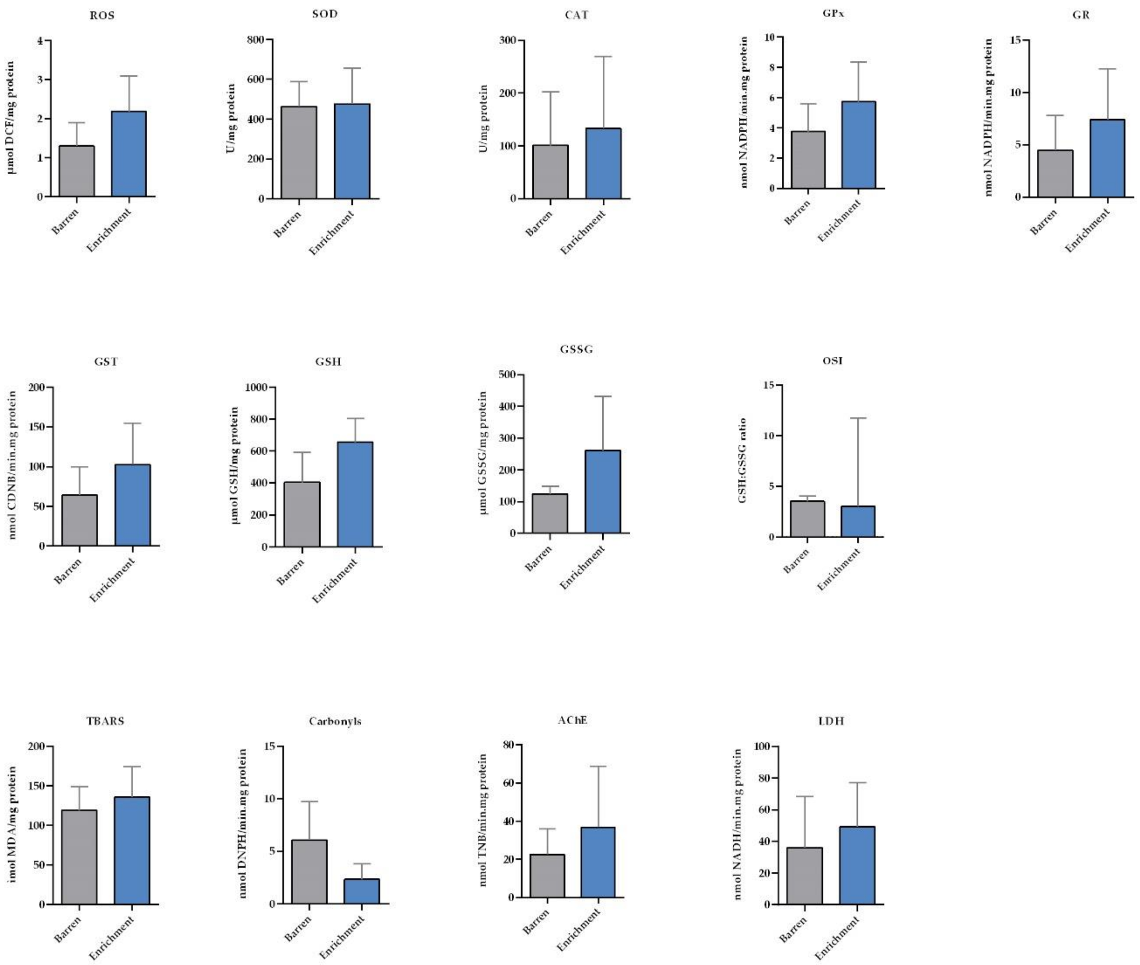

### Table S1

**Table S1.** Significant data ($p < 0.05$) of the random factor Batch from the novel tank test that are not represented in the text or figure. There were no interactions with the fixed factor treatment.

| Variable of the NT | Factor | Description | Statistic test | *p*-value | Statistical test value[1] | Median [IQR] or Mean ± SD |
|---|---|---|---|---|---|---|
| Distance (m) | Batch | First vs Second batch | Univariate analysis | 0.004 | Z= 27424.420 | First:20.24 ± 3.61<br>Second: 13.78 ± 1.82 |
| Average speed (m/s) | Batch | First vs Second batch | Univariate analysis | 0.010 | Z= 4447.864 | First:0.06 ± 0.01<br>Second: 0.04 ± 0.01 |
| Distance (m) during erratic movements | Batch | First vs Second batch | Univariate analysis | 0.026 | Z=603.722 | First:14.75 ± 3.06<br>Second: 4.31 ± 1.98 |
| Duration (s) of the erratic movements | Batch | First vs Second batch | Univariate analysis | 0.037 | Z=296.714 | First:170.83 ± 39.10<br>Second: 67.65 ± 24.57 |
| Distance (m) swam in the BTM zone | Batch | First vs Second batch | Univariate analysis | 0.043 | Z=221.917 | First:17.43 ± 2.01<br>Second: 7.46 ± 1.06 |
| Average speed (m/s) of enriched-housed fish in the UP zone | Batch | First vs Second batch | Independent Samples T Test | 0.022 | t(3)= 4.418 | First:0.08 ± 0.004<br>Second: 0.04 ± 0.007 |

[1] For Independent Samples T test t(df)= t value. For Univariate analysis is presented the standard score (Z); UP (upper) and BTM (bottom) zones of the tank; NT: Novel tank test.
